# The role of intraspecific crop-weed hybridization in the evolution of weediness and invasiveness: cultivated and weedy radish (*Raphanus sativus* L.) as a case study

**DOI:** 10.1101/2023.03.01.530677

**Authors:** Román B. Vercellino, Fernando Hernández, Alejandro Presotto

## Abstract

**PREMISE:** Crop-wild/weed hybrids usually exhibit intermediate and maladapted phenotypes compared to their parents; however, hybridization has sometimes been associated with increased fitness, potentially leading to enhanced weediness and invasiveness. Since the ecological context and maternal genetic effects may affect hybrid fitness, they could influence the evolutionary outcomes of hybridization. Here, we evaluated the performance of first-generation crop-weed hybrids of *Raphanus sativus* L. and their parents under two contrasting ecological conditions.

**METHODS:** Using experimental hybridization and common garden experiments under field conditions, we assessed the differences in time to flowering, survival to maturity, plant biomass and reproductive components between bidirectional crop-weed hybrids and their parents, under two contrasting ecological conditions, agrestal (wheat cultivation, fertilization, weeding) and ruderal (human-disturbed uncultivated area) over two years.

**RESULTS:** Crop, weeds and bidirectional hybrids overlapped -at least partially-during the flowering period, indicating a high probability of gene flow. Hybrids survived to maturity at rates at least as successful as their parents and showed higher plant biomass and fecundity, which resulted in higher fitness compared to their parents in both contrasting environments, without any differences associated with the direction of the hybridization.

**CONCLUSIONS:** Intraspecific crop-weed hybridization, regardless of the cross direction, has the potential to promote weediness in weedy *R. sativus* both in agrestal and ruderal environments, increasing the chances of the introgression of crop alleles into weed populations. This is the first report of intraspecific crop-weed hybridization in *R. sativus*.

## INTRODUCTION

Hybridization is increasingly recognized as an important evolutionary phenomenon that can influence the evolution of both weeds and invasives (Ellstrand et al., 2010, 2013; Vercellino et al., 2023). When hybridization occurs, the hybrid lineages harbor alleles from both progenitors, and the process of introgression will depend on the fitness of the hybrids relative to their parents (Mercer et al., 2007; Presotto et al., 2019). Hybrids commonly exhibit intermediate and maladapted phenotypes and reduced fitness compared to their parents (Mercer et al., 2006; Sahoo et al., 2010; Hooftman et al., 2015; Martin et al., 2019; Yue et al., 2021), with nonviable or sterile offspring in extreme cases (Rieseberg et al., 2007). If novel alleles reduce hybrid fitness, these alleles may not prevail or introgress. However, hybridization can also lead to adaptive evolution through evolutionary novelty, increased genetic variation, hybrid vigor (i.e., heterosis) and dumping genetic load, increasing weediness and invasiveness (Schierenbeck and Ellstrand, 2009; Hovick and Whitney, 2014). Spontaneous hybridization between crop and their wild/weedy relatives has increased the weediness and invasiveness of existing weeds and invasives, e.g., weedy *Brassica rapa* (Ureta et al., 2017; Pandolfo et al., 2018b) and California wild radish (Snow et al., 2010), and has resulted in the evolution of new weeds and invasives, e.g., European weed beet (Bartsch, 2010), weedy rice (Vigueira et al., 2019), European teosinte (Le Corre et al., 2020) and weedy sunflower (Presotto et al., 2017; Hernández et al., 2022). In addition, in some cases, hybrid populations have replaced their progenitors locally (Hegde et al., 2006). Conversely, in many instances, hybridization has not affected weediness or invasiveness (Magnussen and Hauser, 2007; Whitney et al., 2009; Martin et al., 2019; Page et al., 2019).

Hybrid performance relative to their parents may be influenced by the environment (Campbell and Snow, 2007; Arnold and Martin, 2010; Hovick and Whitney, 2014; Presotto et al., 2019). Due to this environmental dependence, evolutionary outcomes of hybridization in agrestal (i.e., agricultural) environments may differ from those in ruderal (human-modified uncultivated area) habitats (Mercer et al., 2006; Presotto et al., 2019; Mitchell et al., 2021). The fitness of wild/weeds could be enhanced by the acquisition of crop traits under agricultural environments. This is especially true for traits conferring resistance to herbicides, insects or diseases, or tolerance to abiotic stresses (Xia et al., 2016; Ureta et al., 2017; Pandolfo et al., 2018b; Nam et al., 2020; Kreiner et al., 2022), but also for other typical crop traits, such as early flowering, high growth rate and seed production (Campbell and Snow, 2007; Presotto et al., 2017; Le Corre et al., 2020). However, the acquisition of crop traits which have been shaped to suit the agricultural environment by artificial selection, frequently create maladapted crop-wild/weed hybrids with lower fitness under less selective environments (e.g., ruderal environments), limiting the adaptive evolution of weeds and invasives (Mercer et al., 2007; Hovick et al., 2012; Presotto et al., 2019; Vercellino et al., 2021a).

Hybrid performance may also be influenced by the phenotypes of its maternal parent (known as maternal genetic effects) (Kimball et al., 2008; Hernández et al., 2021). Maternal genetic effects (hereafter maternal effects) are defined as the contribution of the phenotype or genotype of the maternal parent to the phenotype of its offspring beyond the equal chromosomal contribution expected from each parent (Roach and Wulff, 1987). Numerous studies have investigated the ecological and evolutionary consequences of crop-to-wild/weed hybridization (Campbell et al., 2006; Mercer et al., 2006, 2007; Snow et al., 2010; Hovick et al., 2012; Presotto et al., 2019). However, much less attention has been paid to hybridization in the opposite direction (i.e., wild/weed-to-crop hybridization) and how maternal genetic effects may influence the fitness of their offspring immediately after hybridization (Kimball et al., 2008; Piskurewicz et al., 2016; Singh et al., 2017; Hernández et al., 2021). If such maternal effects exist, crop-weed hybrids produced on crop and weed plants will show differences in performance. Common garden studies comparing the fitness of bidirectional crop–wild/weed hybrids relative to both parents under contrasting ecological conditions are essential to deepen our understanding on the role of crop-wild/weed hybridization on weediness and invasiveness (Mercer et al., 2006; Campbell and Snow, 2007).

In this study, we performed common garden experiments using experimental populations of the crop-weed complex of radish (*Raphanus sativus* L.) as a model system to assess the hypothesis that bidirectional intraspecific crop-weed hybridization increases weediness and invasiveness (Bourgeois et al., 2019). These studies allow the acceleration of natural processes in order to explore local ecological scenarios under which the fitness of crop-weed hybrids may equal or exceed the fitness of their parents, favoring the persistence of crop alleles in wild/weedy populations and consequently enhancing weediness and invasiveness (Mercer et al., 2006; Campbell et al., 2009). The *Raphanus* complex, formed by the cultivated species *R. sativus*, their weedy conspecific forms and their weed related species *R. raphanistrum*, is a well-established model system that has been widely used to evaluate the ecological and evolutionary consequences of interspecific crop-weed hybridization (Campbell et al., 2006, 2009; Hegde et al., 2006; Campbell and Snow, 2007; Ridley et al., 2008; Snow et al., 2010; Hovick et al., 2012; Shukla et al., 2020, 2021). We compared the time to flowering, survival to maturity, plant biomass and reproductive components of bidirectional first-generation crop-weed hybrids in relation to both parents under two contrasting ecological environments, agrestal (wheat cultivation, fertilization, weeding) and ruderal (human-disturbed uncultivated area) over two years. Despite that the fitness of first-generation hybrids does not determine the ultimate fate of all crop alleles, it affects the initial course of introgression through its effect on the frequencies of crop alleles that are subsequently subject to selection and drift (Snow et al., 1998; Mercer et al., 2006; Presotto et al., 2019). Our work represents the first experimental study to evaluate the effect of bidirectional crop-weed hybridization in *R. sativus* on the evolution of weediness and invasiveness.

## 2. MATERIALS AND METHODS

### Study system

Conspecific cultivated and weedy radish (*Raphanus sativus* L., Brassicaceae) are annual, self-incompatible, insect-pollinated plants, that can hybridize over long distances (Ellstrand et al., 1989; Snow and Campbell, 2005). Cultivated radish is an ancient crop domesticated independently in both Europe and South Asia (Zhang et al., 2021) and selected for multiple purposes, including human consumption as roots (botanically hypocotyl plus root), forage and cover crops (Snow and Campbell, 2005). *R. sativus* is not known in the wild (Warwick, 2011); however, spontaneous populations of radish have been found as a weed infesting crops in several parts of the world, including North America, South America and South Africa (Hegde et al., 2006; Barnaud et al., 2013; Vercellino et al., 2018; Costa et al., 2021; Heap, 2023). In North America, its invasive capacity is attributed to the introgression with a closely related species *R. raphanistrum* (known as California wild radish) (Hegde et al., 2006; Ellstrand et al., 2010), and in South America, it can be found in Argentina, Brazil, Chile, Uruguay and Paraguay (Pandolfo et al., 2018a). In Argentina, weedy radish is present in 20 out of 23 provinces and has been considered a widespread invasive weed since at least the 1930s (Ibarra, 1937; Pandolfo et al., 2018a). It grows in disturbed environments, such as road margins, rural roads and fence lines, and within agricultural fields. It is one of the most noxious weeds in cereals, oilseeds and some horticultural and forage crops (Scursoni et al., 2014; Vercellino et al., 2018, 2021b) and has evolved resistance to AHAS (acetohydroxyacid synthase, also known as acetolactate synthase (ALS))-inhibiting herbicides (Pandolfo et al., 2016; Heap, 2023). Argentine weedy *R. sativus* populations show many crop-like traits, such as white or purple flowers, pods attached to the mother plant at maturity, and no seed dormancy (Vercellino et al., 2018, 2019). On the other hand, weedy populations do not harbor typical traits of *R. raphanistrum* like constricted fruits and yellow flowers (Pandolfo et al., 2018a). However, they have an extended period of seedling emergence due to the requirement of darkness for germination and the mechanical restriction of the pericarp (Vercellino et al., 2019).

In southern Buenos Aires, an area severally infested with weedy radish (Scursoni et al., 2014; Vercellino et al., 2018), the use of cultivated radish as horticultural and cover crop has grown rapidly in recent years, leading to the formation of natural crop-weed hybrids in the field.

### Germplasm and crosses

In this study, bidirectional crosses between weed and cultivated *R. sativus* were used due to the potential bidirectional hybridization in the *Raphanus* complex in Argentina. Weed populations were collected from two sites in southern Buenos Aires: Balcarce (BAL; 37°35’25”S, 58°31’59”W) in 2008 and Pieres (PIE; 38°24’31”S, 58°35’60”W) in 2011 (Vercellino et al., 2018), before the introduction of *R. sativus* cover crop cultivars in those areas. Balcarce and Pieres are temperate sub-humid environments, with an annual average temperature of 13.9 and 14.3 °C (SMN 2023, https://www.smn.gob.ar/), respectively. Cultivated materials were represented by two commercial cultivars: the European radish ‘Rovi – Round Red Radish with White Tips’ (Garde, Giusti y Chuchuy S.A., Ciudad Autónoma de Buenos Aires, Argentina) cultivated for root consumption and the daikon radish ‘CCS 770 – Daikon Radish’ (El Cencerro, Coronel Suarez, Buenos Aires, Argentina) cultivated as cover crop. Radish is cultivated in urban and peri-urban farms and gardens throughout the country, and as cover crop radish in the Pampas region and northern Argentina, Paraguay and Brazil (Snow and Campbell, 2005). Bidirectional crosses were made between weed populations and cultivars (referred to as CROP). The crosses (hereafter referred to as Biotypes) were characterized according to their female and male parents.

To minimize the maternal environmental effects, all the seeds used in each experiment were produced in a greenhouse (20 ± 5°C, natural light) in the Agronomy Department at Universidad Nacional del Sur, Bahía Blanca, Argentina. For seed production, 60 plants of each BAL, PIE, and CROP were grown in the greenhouse and seeds were produced under controlled hand-pollination, creating seven different biotypes: three pure parents (BAL, PIE and CROP) and bidirectional F1 crop-weed hybrids (BAL x CROP, PIE x CROP, CROP x BAL and CROP x PIE). CROP was represented by the open-pollinated cultivar ‘Rovi – Round Red Radish with White Tips’ in 2019, and by the cultivar ‘CCS 779 Daikon Radish’ in 2020. In the greenhouse, the weedy and cultivated plants flowered simultaneously. Seeds were produced as follows: at the flowering stage, unopened flower buds from weed plants were opened and emasculated without damaging the pistil by using sharp-edged forceps before pollen dehiscence. Once emasculated, the flowers were hand-pollinated with pollen from sibling plants (weed biotypes: BAL and PIE) or pollen from cultivated plants (weed-crop hybrids: BAL x CROP and PIE x CROP). Likewise, unopened flowers buds from cultivated plants were opened, emasculated and hand-pollinated with pollen from sibling plants (CROP biotype) or pollen from weed plants (crop-weed hybrids: CROP x BAL and CROP x PIE). Prior to emasculation, all other flower buds on the racemes were removed. Immediately after pollination, the racemes were covered with 12 × 6 cm pollination bags to avoid any pollination from other plants. Pod formation from successful hand-pollination could be observed ten days after pollination and, subsequently, the bags were removed from the racemes. The effectiveness of pollination was >99 %. At the end of the growing season, the mature pods were stored at room temperature and less than 10% moisture until use. For experiments in the 2020 and 2021, seeds were produced during the 2019 and 2020 growing seasons, respectively.

### Site and experimental design

We compared weed, crop, and their hybrids in two environments (ruderal and agrestal). Common garden experiments were conducted in the Agronomy Department experimental field at the Universidad Nacional del Sur, Bahía Blanca, Argentina (38°41´38” S, 62°14´53” W), during two winter growing seasons (Jun-Jan 2020/21 and 2021/22, the typical life cycle for both the crop and weedy radish). The site is characterized as semi-arid with a temperate climate, and a mean annual temperature of 14.9 °C; the mean temperatures of the coldest month (July) and the hottest month (January) are 7.6 and 23.0 °C, respectively. The average annual precipitation is 641 mm (SMN 2023, https://www.smn.gob.ar/). The soil has a well-drained loamy sand texture with 1.1% organic matter, pH 7.7, and 8.4 mg kg^−1^ (0–12 cm) available phosphorus (P) (a typical soil in the southwest of Buenos Aires province). The experimental field was maintained as a natural field for at least two years prior to the start of the experiment, leaving natural vegetation to grow. The site was governed by herbaceous plants, mainly *Lolium spp.* and *Diplotaxis tenuifolia* in winter, and C*entaurea solstitialis, Chenopodium album, Portulaca oleracea, Helianthus annuuss* subsp*. annuus, Cynodon dactilon* and *Cenchrus incertus* in spring-early summer. One month prior to wheat sowing (see below), a tillage operation with a disc harrow was used to prepare the soil. Then the environments were defined. Field experiments were arranged in a split-plot design. Environments (agrestal and ruderal) were assigned to main plots (one main plot per environment, that is in total two main plots). Each main plot was divided to eight blocks, and each block was divided in eight subplots. The seven biotypes (CROP, BAL, PIE, CROP x BAL, CROP x PIE, BAL x CROP and PIE x CROP) plus empty control were assigned to subplots within each block, whereby the assignment to subplot was randomized for each block, environment and year. Each subplot contained ten plants of the same biotype (Appendix S1; see the Supplementary Data with this article). The whole experiment was repeated in two subsequent years.

**Appendix S1.**
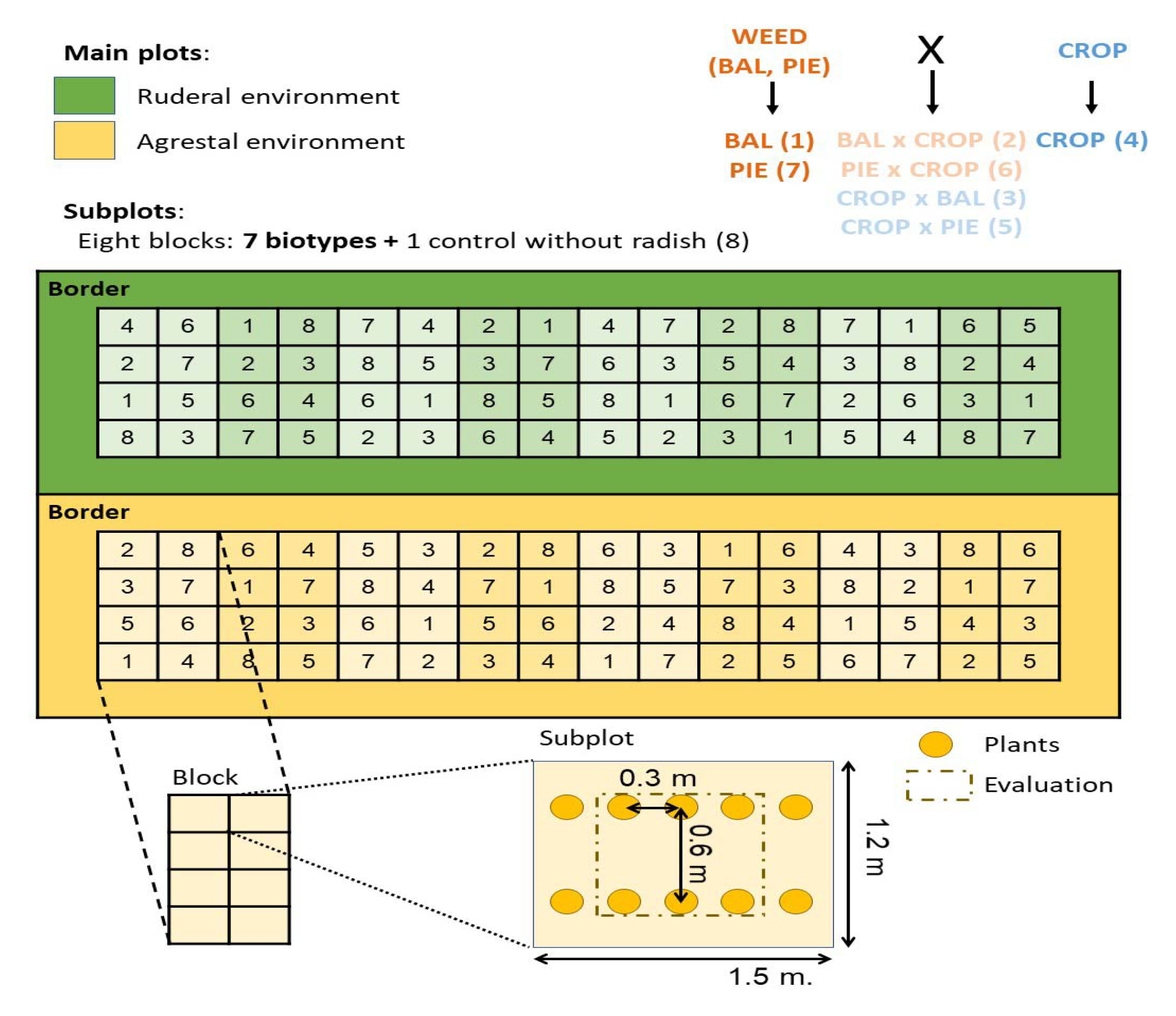
Experimental crosses and common garden design. Environments (main plots) are indicated in different colors, agrestal (yellow) and ruderal (green). Each environment was composed of eight blocks, each of eight subplots. Subplots were seven radish biotypes (1- BAL, 2-BAL x CROP, 3-CROP x BAL, 4-CROP, 5-CROP x PIE, 6-PIE x CROP, 7-PIE) and a control plot without radish (8). Numbers (1-8) on the field map represent each biotype or control position in year 1. In year 2, both environments were randomized again.

From the radish seeds produced in the crossing design described above, 2-3 seeds per pod from more than 400 pods were pooled to represent each biotype. Since seeds from both weed and cultivated radish are non-dormant (Vercellino et al., 2019), radish seeds from all seven biotypes were hand-planted in early Jul 2020, 2-3 seeds per planting positions, 2-3 cm deep, at 0.3-m intervals in rows spaced at 0.6-m. The subplots were two rows wide by five planting positions long. Sixty-four subplots ((7 biotypes + 1 control without radish) x 8 blocks) made up each main plot (i.e., each ruderal and agrestal environment). The main plots were surrounded by a border of two rows of weedy radish to buffer against possible border effects. Since most of the seeds germinated and emerged, at the fully expanded one-leaf stage, radish seedlings were hand-thinned to one per position, reaching the target of 10 plants per subplot. In 2021, since the experiment was established in a place where there was weedy radish three years before and to avoid contamination from the seed bank, seedlings of the seven radish biotypes were established in plastic trays containing potting mix (Grow Mix Terrafertil®), grown in a glasshouse (20 ± 5°C, natural light and automatic watering), and transplanted into the experimental field at the 2-3-leaf stage, in early Aug, as above. Experiments were drip irrigated, watered complementarily at the beginning to facilitate their establishment, and later when necessary.

### Description of environments

#### Agrestal environment

Before the establishment of the radish biotypes in the field in late autumn (mid-Jun), in the agrestal environment, a wheat (*Triticum aestivum* L.) crop was established as follow: seeds were planted in 0.2-m spaced rows at 3-cm deep, to reach a target of 160 plants m^-2^ (the common stand density in southern Buenos Aires). Wheat rows were arranged to surround rather than overlap the radish plants. The cultivars of wheat used were ACA 360 and KLEIN MINERVA in 2020/21 and 2021/22, respectively. Fertilization consisted of 120 kg ha^-1^ diammonium phosphate at planting, 125 kg ha^-1^ urea at early tillering and 125 kg ha^-1^ urea at mid-tillering. Weeds other than radish were periodically removed by hand. At the end of each experiment, the yield of the wheat crop was estimated by hand-harvesting and hand-threshing four replicates of 1 m^2^, collected in subplots without radish. The yield of the wheat was 5136 ± 201 and 5190 ± 188 kg ha^-1^ in 2020/21 and 2021/22, respectively.

#### Ruderal environment

In the ruderal environment, we did not establish any crops or fertilize the soil. Biotypes were exposed to naturally occurring weather conditions, competing weeds (i.e., same as described above), herbivores, and pathogens, simulating a human-disturbed uncultivated area (e.g., field margin, fence row).

### Measurements

In all ten plants per subplot, we recorded daily time to flowering (days from emergence to first flowering) and survival to reproduction (defined as plants that produced at least one pod). At maturity, the above-ground biomass of the six central radish plants from each subplot was hand-harvested, dried at 60°C to constant weight and weighed to determine the total aboveground dry biomass per plant (hereafter referred to as ‘plant biomass’). After that, pod biomass per plant, pod number per plant, seed biomass per plant, seed number per plant and seed weight were evaluated in the harvested plants following the procedure described in Vercellino et al. (2018, 2021b). Seed number per plant was used as the proxy of fecundity.

Since all ten plants per subplot were used to evaluate survival to maturity, but only the six central plants to estimate fecundity, the mean fitness of each subplot was estimated by multiplying survival to maturity by the average fecundity of each subplot in order to calculate the fitness of bidirectional crop-weed hybrids relative to weed plants (hereafter referred to as ‘relative fitness’). Relative fitness (RF) was calculated as: RF = bidirectional hybrid (crop-weed or weed-crop)/weed for each block and weed origin (BAL and PIE).

### Statistical Analysis

All data were analyzed using generalized linear mixed models (GLMMs) with restricted maximum likelihood in PROC GLIMMIX (SAS University Edition). We analyze the effects of experimental factors in the time to flowering, survival to reproduction, total aboveground biomass, and reproductive components (i.e., pod biomass per plant, pod number per plant, seed biomass per plant, seed number per plant and seed weight). For each year (2020/21 and 2021/22), environment (ruderal and agrestal), biotype (CROP, BAL, PIE, CROP x BAL, CROP x PIE, BAL x CROP and PIE x CROP), and its interaction (environment x biotype) were considered as fixed effects, while the block nested within environment effect was considered as random. After that, we grouped the seven biotypes into four groups: WEED (BAL and PIE), WEED x CROP (crop-weed hybrids produced on weed plants: BAL x CROP and PIE x CROP), CROP x WEED (crop-weed hybrids produced on crop plants: CROP x BAL and CROP x PIE), and CROP. To compare groups (WEED, WEED x CROP, CROP x WEED, and CROP), we constructed six a priori orthogonal contrasts: (1) WEED vs. CROP x WEED (2) WEED vs. WEED x CROP, (3) WEED vs. CROP, (4) CROP x WEED vs. WEED x CROP, (5) CROP x WEED vs. CROP, and (6) WEED x CROP vs. CROP, and compare them using the CONTRAST statement in PROC GLIMMIX, SAS.

Since categorical analyses do not allow for the complex model specifications needed in an analysis of a split-plot design, survival to reproduction was analyzed as a continuous response variable, based on averages of radish plants in each subplot. Data were arcsine-square-root transformed prior to analyses to meet assumptions of normality (Mercer et al., 2006). Flowering time analysis only included radish plants that reached flowering, and plant biomass and reproductive component (pod biomass per plant, pod number per plant, seed biomass per plant, seed number per plant and seed weight) analyses only included radish plants that survived to reproduction. To avoid redundancy, we performed a multiple Pearsońs correlation analysis between plant biomass and the reproductive components with PROC CORR in SAS, and for all pairwise comparisons with r > |0.9|, and we only retained one variable of each pair. After this filtering, we retained seed number per plant and seed weight (Appendix S2). All traits (i.e., except survival to reproduction which was previously indicated) were square-root transformed to satisfy assumptions regarding the normality of the residuals. Finally, least square means of relative fitness were estimated using PROC GLIMMIX.

**Appendix S2.** Correlations among plant biomass and reproductive components of weedy radish.

| Traits | Biomass<br>plant<sup>-1</sup> | Pod<br>biomass<br>plant<sup>-1</sup> | Pod<br>number<br>plant<sup>-1</sup> | Seed<br>biomass<br>plant<sup>-1</sup> | Seed<br>weight | Seed number<br>plant<sup>-1</sup> |
| --- | --- | --- | --- | --- | --- | --- |
| Biomass plant<sup>-1</sup> | 1 | **0.986** | **0.942** | **0.958** | 0.074 | **0.952** |
| Pod biomass plant<sup>-1</sup> |  | 1 | **0.943** | **0.972** | 0.092 | **0.957** |
| Pod number plant<sup>-1</sup> |  |  | 1 | **0.921** | 0.015 | **0.934** |
| Seed biomass plant<sup>-1</sup> |  |  |  | 1 | 0.088 | **0.979** |
| Seed weight |  |  |  |  | 1 | -0.009 |
| Seed number plant<sup>-1</sup> |  |  |  |  |  | 1 |

Correlation coefficients are reported. The pairwise correlations with $r > |0.9|$ are in bold. N> 1000 for each combination.

## RESULTS

### Time to flowering

In year 1, there were significant biotype, environment, and biotype x environment interaction effects; therefore, we performed contrasts for each environment (agrestal and ruderal) (Table 1). In both environments, the crop flowered later than the weeds, although differences were larger in the ruderal environment (Fig. 1a, Table 1). Overall, hybrids showed a flowering time intermediate to that of both parents. Whereas the crop-weed hybrids produced on weed plants flowered simultaneously with the weeds, the crop-weed hybrids produced on crop plants flowered a little bit later (<5 days) than the weeds, resulting in significant differences between bidirectional hybrids only in the agrestal environment (Fig. 1a, Table 1). In year 2, there were significant biotype and environment effects, but no biotype x environment interaction (Table 2). Crop, weeds and hybrids flowered simultaneously (Fig. 1b), although some contrasts showed statistically significant (<3 days) differences between groups (Fig. 1b, Table 2).

**Figure 1.**
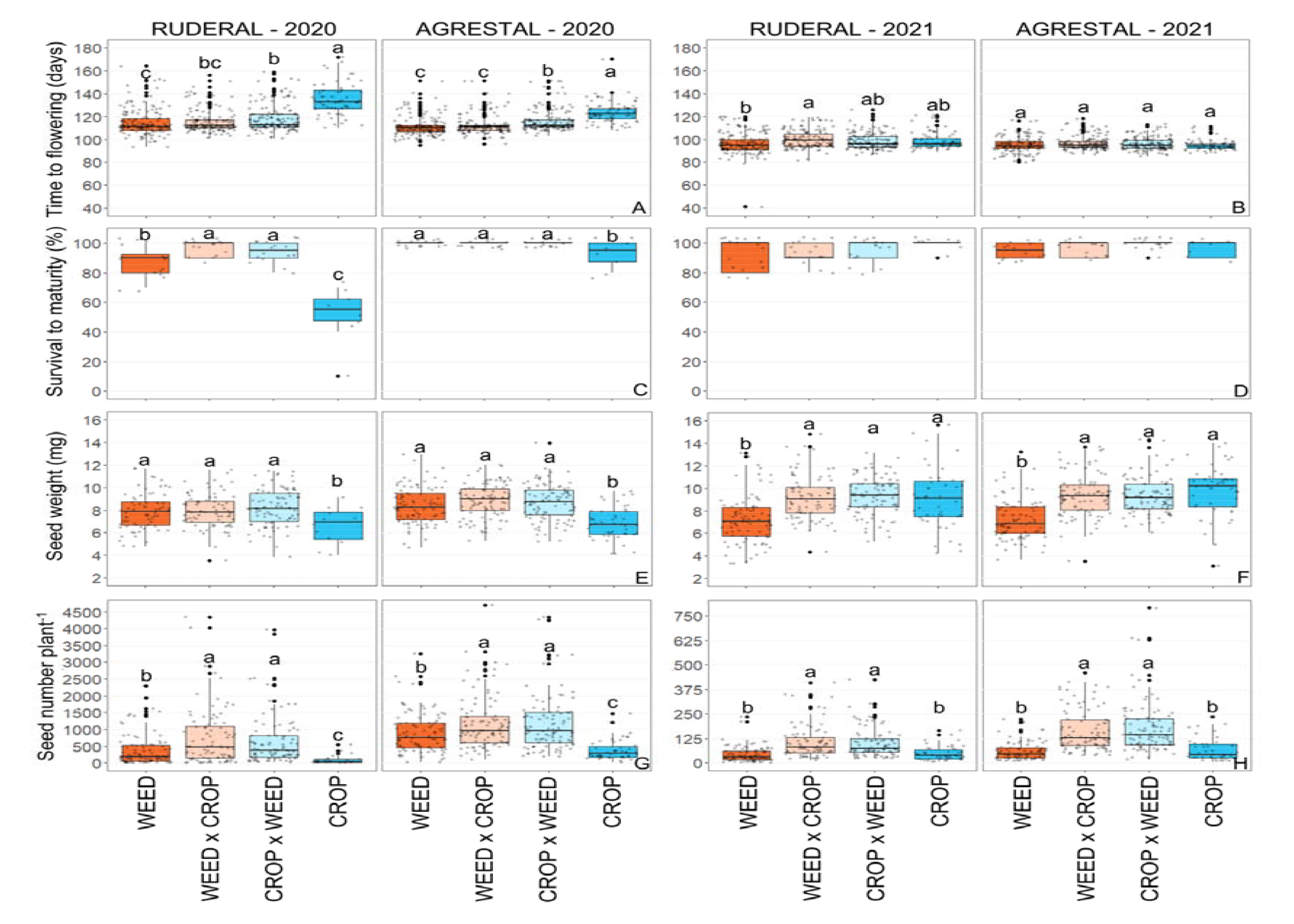
Boxplots of time to flowering (2020: A; 2021: B), survival to reproduction (2020: C; 2021: D), seed weight (2020/21: E; 2021/22: F) and seed number per plant (2020: G; 2021: H) for bidirectional crop-weed hybrids and their parents (WEED: BAL and PIE, CROP) of *Raphanus sativus* growing in two contrasting environments (ruderal and agrestal), and in two years. Box edges represent the 0.25 and 0.75 quartiles, solid line represents the median value, and whiskers extend to the minimum and maximum value or 1.5x the interquartile range with points for more extreme values. Black points represent outliers, and small and transparent points represent each data point. Values sharing the same letter within an environment and year are not significantly different based on contrast tests, and different letters within a panel indicate significant differences as generated by the contrasts. Note the variation in the scale of the y-axis between (H) and (G) to show large differences in seed number per plant between year 1 and 2.

**Table 1.**
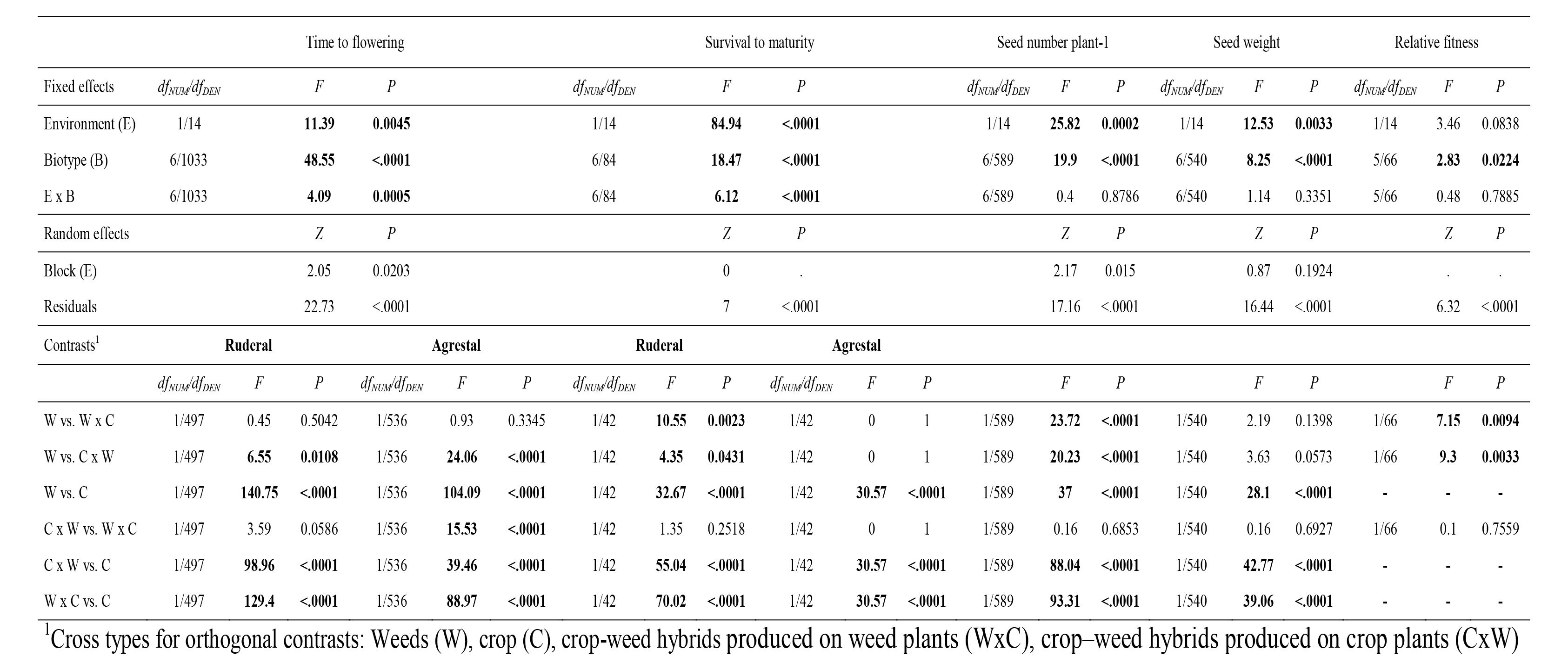
Restricted maximum likelihood for survival to reproduction, days to flowering, seed weight and seed number per plant in crop-weed bidirectional hybrids and their progenitors (weed: BAL and PIE; CROP) in two contrasted environments (ruderal and agrestal) performed using maximum likelihood in SAS PROC GLIMMIX for year 1. Data were collected in the Agronomy Department at Universidad Nacional del Sur, Bahía Blanca, Argentina.

**Table 2.**
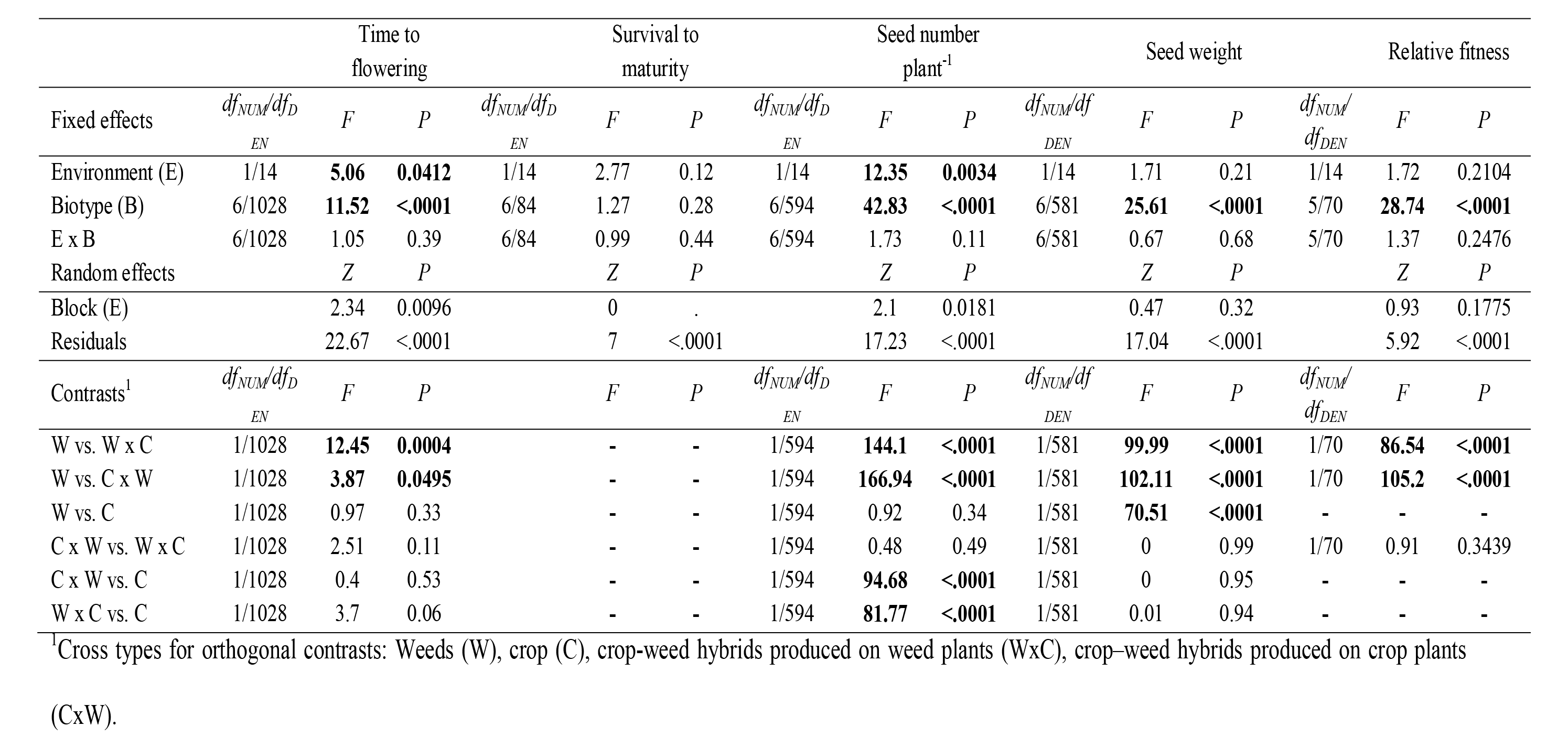
Restricted maximum likelihood for survival to reproduction, days to flowering, seed weight and seed number per plant in crop-weed bidirectional hybrids and their progenitors (weed: BAL and PIE; CROP) in two contrasted environments (ruderal and agrestal) performed using maximum likelihood in SAS PROC GLIMMIX for year 2. Data were collected in the Agronomy Department at Universidad Nacional del Sur, Bahía Blanca, Argentina.

### Survival to maturity

In year 1, there were significant biotype, environment, and biotype x environment interaction effects; therefore, we performed contrasts for each environment (agrestal and ruderal) (Table 1). In the agrestal environment, all the weed and hybrid plants survived to maturity, whereas the crop survived significantly less than the weeds and hybrids (Figure 1c, Table 1). In contrast, in the ruderal environment, there was more variation between groups; hybrids (both cross directions) survived significantly more than weeds, and weeds survived significantly more than the crop (Figure 1c, Table 1). There were no significant differences between bidirectional hybrids (Figure 1a, Table 1). In year 2, most plants (> 95 %) survived to maturity and there were no significant effects of environments, biotypes, nor environment x biotype interaction (Figure 1b, Table 2).

### Fecundity: reproductive components

For the set of six variables (plant biomass, pod biomass per plant, pod number per plant, seed biomass per plant, seed number per plant and seed weight), we found positive correlations between seed number per plant and plant biomass (r = 0.95), pod number per plant (r = 0.93), pod biomass per plant (r = 0.96) and seed biomass per plant (r = 0.98), while the seed weight showed no correlation (r < 0.10) with any of these traits (Appendix S2). Therefore, seed number per plant and seed weight were retained for further analysis.

Analysis of the seed number per plant (i.e., fecundity) revealed significant biotype and environment effects, but no significant biotype x environment interaction in either year (Tables 1 and 2). Overall, all plants in the agrestal environment showed 84.3 and 60.3 % higher fecundity than the radish plants in the ruderal environment in year 1 and 2, respectively (Figs. 1g-h, Tables 1 and 2). Fecundity of weeds was 167.5 % higher than crop in year 1, but no differences were observed in year 2 (Figs. 1g-h, Tables 1 and 2). Fecundity of the crop-weed hybrids were 47.0 and 169.0 % higher than fecundity of the weeds in year 1 and 2, respectively, with no differences associated with the cross direction, indicating no maternal genetic effect (Figs. g-h, Table 1).

With respect to the seed weight, there were significant biotype and environment (only in year 1) effects, and no significant biotype x environment interaction (Tables 1 and 2). In year 1, the seeds of the weed plants were 21.3 % heavier than the seeds of the crop plants. However, in year 2, the seeds of the crop plants were 33.4 % heavier than the seeds of the weed plants (Figs. 1e-f, Tables 1 and 2). In both years, the seeds of crop-weed hybrids (regardless of the cross direction) were as heavy as the seeds of the parent with heavier seeds (Figs. 1e-f, Tables 1 and 2).

### Relative fitness

In both years, analysis of the relative fitness revealed a significant biotype effect but no significant environment or biotype x environment interaction effects (Tables 1 and 2). The relative fitness of hybrids compared to weeds was 1.86 ± 0.21 and 3.11 ± 0.20 in year 1 and 2, respectively, regardless of the environment and the direction of hybridization (Tables 1 and 2).

## DISCUSSION

This study shows that crop-weed hybridization has the potential to stimulate weediness in weedy *R. sativus* under both agrestal and ruderal environments. First-generation crop-weed hybrids showed hybrid vigor in all measured traits; they survived at rates that are at least as successful as the rates of both weed and crop plants and produced higher plant biomass that resulted in 47 and 169 % more fecundity than the weeds in year 1 and 2, respectively, in both ruderal and agrestal environments. Relative fitness of hybrids compared to weeds was 1.86 and 3.11 in year 1 and 2, respectively, with no differences between environments. Seeds of hybrids were as heavy as the seeds of the parental biotype with heavier seeds. Finally, we found no differences associated with the direction of hybridization, indicating the general absence of maternal genetic effects, which also supports hybrid vigor as the main genetic mechanisms of increased fitness. Our study highlights the risk of increasing weediness after hybridization in *R. sativus*, and suggests hybrid vigor as the genetic mechanisms increasing fitness of hybrid progeny.

### Evolutionary consequences of crop-weed hybridization

In our experiments, cultivated radish started flowering later than the weedy radish in year 1, but both cultivated and weedy radish started flowering simultaneously in year 2. This is probably associated with genetic differences between the radish cultivars. In year 1, we expected that the crop-weed hybrids would have delayed flowering compared to weedy radish, as was observed in California wild radish (hybrids between *R. sativus* and *R. raphanistrum*) (Campbell et al., 2006, 2009). While we found a trend in this direction (especially in the crop-weed hybrids produced on crop plants) (Fig. 1a), the difference was less than five days. Due to the long flowering period of the species, the flowering period of cultivated and weedy radish largely overlapped in both years in the field and in the greenhouse during seed production (data not shown), suggesting that crop-weed hybridization is highly probable if crop and weed grow in sympatry.

Common garden studies using experimental early generation crop-weed hybrids can provide tentative predictions of the effects of hybridization on the dynamics of weed populations and the likelihood of crop allele introgression into weeds (Campbell et al., 2006; Mercer et al., 2006; Presotto et al., 2019). These studies allowed us examining effects of crop-weed hybridization on early and later life history traits over environmental conditions that realistically represent those at interfaces between crop and weed populations (Mercer et al., 2006; Presotto et al., 2019). In contrast to the common assumption (based on previous results) that early generations of crop-wild/weed hybrids suffer dramatic reductions in fitness relative to their wild/weedy counterpart (Snow et al., 1998; Cummings et al., 2002; Mercer et al., 2006; Campbell and Snow, 2007; Sahoo et al., 2010; Hooftman et al., 2015), our results showed at least equal survival and vastly higher plant biomass and fecundity in the hybrids relative to their parents in both contrasting environments and years. The increased fitness of first-generation hybrids is expected to facilitate the introgression of crop alleles into weedy populations and stimulate the evolution of weediness (Schierenbeck and Ellstrand, 2009; Vercellino et al., 2023). Similarly, potential for introgression from cultivated to their wild or weedy counterpart has been found in other similar crop-wild/weedy complexes, such as wild beet (Bartsch, 2010), wild lettuce (Uwimana et al., 2012), wild chicory (Kiær et al., 2007), wild carrot (Magnussen and Hauser, 2007), weedy *Brassica rapa* (Hauser et al., 2003) and wild radish (Hegde et al., 2006).

The assessment of the effect of crop-weed hybridization under contrasting ecological conditions was used to evaluate the environmental dependence of hybridization effects. In previous studies, the weed *R. raphanistrum* showed higher fecundity than the crop-weed hybrids under non-competitive conditions, but the crop-weed hybrids were less negatively affected under intense competition, suggesting that hybrids may be favored under the highly competitive condition imposed by crops in agricultural environments (Campbell and Snow, 2007; Campbell et al., 2009). Similarly, wild sunflower (*Helianthus annuus*) showed higher fecundity than crop-wild hybrids in four contrasting environments, but the relative fitness of hybrids was higher under competitive conditions, possibly because wild sunflower is not adapted to agricultural environments (Mercer et al., 2006, 2007). Contrary to our expectation, our results showed higher plant biomass and fecundity and consequently higher relative fitness in the hybrids compared to their parents in both contrasting environments, indicating that the result of the crop-weed hybridization was not environmental dependent, and that bidirectional crop-weed hybridization has the potential to increase weediness in a wide range of environmental conditions (e.g., both inside and outside the agricultural environment). However, our inference should be studied further by evaluating the effect of crop-weed hybridization across a greater diversity of locations in order to assess the generality of these results.

Comparison of the direction of crop-weed hybridization was associated with the fact that bidirectional crop-weed hybridization may contribute to the evolution of weediness in weedy radish (Ridley et al., 2008), and previous studies have shown differential maternal genetic effects when evaluating crop-wild/weed hybridization (Pace et al., 2015; Singh et al., 2017; Hernández et al., 2021). Contrary to our expectation, the evaluated traits were not generally influenced by the genotype of the maternal parent, suggesting similar evolutionary consequences regardless of the direction of the hybridization. Nonetheless, considering that radish was selected for larger root and leaf size -unlike crops selected for grain production such as corn, sunflower and rice-and the lack of substantial differences between crop and weedy radish, the absence of maternal genetic effects is not outlandish. To our knowledge, this is the first study that has evaluated plants of bidirectional crop-weed hybrids in a common garden under field conditions.

A plausible explanation for the increased plant biomass and fecundity of hybrids is heterosis (Ellstrand and Schierenbeck, 2000). Heterosis, defined as the superiority of hybrids relative to their parents, is expected to occur after admixture between divergent populations that experienced genetic bottlenecks (Hovick et al., 2012; van Kleunen et al., 2015). Weedy radish populations from Argentina probably experienced strong genetic bottlenecks, first as a result of the species domestication, followed by species introduction into Argentina, and then as result of the feralization process (Vercellino et al., 2023). However, if current cultivars and weedy populations are divergent enough to explain heterosis effects observed in this study is unknown. Genetic studies are needed to better understand the evolutionary history of weedy radish in Argentina. Additional studies using weed-weed and crop-crop hybridization would provide control crosses where there is mixing between populations, but no specific crop-weed hybridization, and help us to disentangle the potential mechanisms of how crop-weed hybridization could lead to hybrid vigor and adaptive evolution. Theory predicts that heterosis of early generation hybrids is only transitory and it would decrease in the following generations (Rius and Darling, 2014); however, previous studies have shown that hybrid vigor could be maintained or fixed in subsequent generations (Ellstrand and Schierenbeck, 2000). For example, the superiority of interspecific crop-weed (*R. sativus* x *R. raphanistrum*) hybrids was maintained at least after four generations and under several environmental conditions (Campbell et al., 2006, 2009; Campbell and Snow, 2007; Snow et al., 2010; Hovick et al., 2012). Similar results were found in natural interspecific crop-weed radish hybrids (Hegde et al., 2006; Ridley and Ellstrand, 2009). However, additional studies should be carried out to understand the long-term consequences of intraspecific hybridization events (Campbell et al., 2009; Snow et al., 2010; Mercer et al., 2014; Mitchell et al., 2019).

A possible limitation of our study of bidirectional crop-weed hybridization is that only two crop cultivars and two weed populations were used to create the biotypes. Despite this, we did not find significant differences in any trait as a result of the sources of the weed population, which could somehow support the generality of our results. On the other hand, hybrid fitness may be influenced by factors that were not considered in our common garden experiments, e.g., seed dormancy and seedling emergence (e.g., Hooftman et al. 2015; Mitchell et al. 2019). However, fecundity is the most important trait contributing to hybrid population growth in the *Raphanus* complex of North America (Campbell et al., 2014, 2016; Teitel et al., 2016). Future studies incorporating more life-history traits would be necessary in order to better understand the result of crop-weed hybridization. Finally, our discussion and conclusions are based on field experiments representing the two main ecological conditions that progenitors originally colonize and where bidirectional hybridization can occur; however, while we adequately tested the effect of crop-weed hybridization within each environment, comparisons between environments within year should be carried out carefully as they were not replicated.

### Implications for crop and weed management

The assessment of the effect of intraspecific crop-weed hybridization under contrasting ecological conditions was initially motivated by concerns about the risk of the evolution of weediness after crop-weed hybridization. Here, we discuss the implications of our results for weed evolution and management.

In recent years, *R. sativus* has been included as a cover crop in agricultural systems from several parts of the world (Lawley et al., 2011). In the Argentine Pampas region, radish cover crop is currently cultivated for both seed production and in farm fields in the same areas where weedy radish has been one of the most problematic weeds for more than 80 years (Ibarra, 1937; Scursoni et al., 2014). Considering that *R. sativus* is a self-incompatible, insect-pollinated species (Snow and Campbell, 2005) with high gene flow over long distances (Ellstrand et al., 1989), crop-weed hybridization is expected in areas where cultivated and weedy radish are sympatric. Therefore, the concern regarding crop-weed hybridization and the subsequent evolution of weediness is not whether it can happen, but the speed with which humans accelerate this evolutionary process (reviewed in Vercellino et al. 2023).

A management practice to prevent crop-weed gene flow could be the suppression of the cultivated radish prior to the flowering stage. Since this practice could only be carried out in farmer fields, but not in seed production areas where the risk of dispersion is considerably higher (e.g., weedy beet in Europe; Bartsch 2010), we propose that the introduction of cultivated radish in areas with weedy radish should be carried out with extreme care, i.e., by planting cultivated radish in weedy radish-free plots and by controlling weedy radish plants in its surroundings. Considering that weed evolution after the introduction of a new crop in a new area has been demonstrated in several crop-weed complexes, with multiple negative impacts (reviewed in Ellstrand et al. 2010; Vercellino et al. 2023), we recommend a thorough agroecological and environmental risk assessment of the introduction of a new crop (e.g., radish cover crop) before being approved for production and commercialization. In addition, integrated weed management strategies aimed at preventing gene flow with the crop should be a priority in order to prevent the emergence and spread of novel more problematic weedy radish biotypes (reviewed in Vercellino et al., 2023).

Finally, crop-weed hybridization not only has negative consequences for maintaining crop genetic uniformity and the increase in weediness, but weedy radish and hybrid plants could also be used as genetic resources to improve crops, including crops more tolerant to biotic and abiotic stress or more locally adapted (Wu et al., 2021; Mabry et al., 2023). In our case, higher biomass and fitness of the hybrids could be interesting for the production of improved cultivated radish both for cover crops or fodder, with higher production of dry matter. However, the risk of fertilization of new cultivars should be taken into account.

## CONCLUSION

In summary, our results using experimental populations under field conditions in a common garden show that crop-weed hybridization has the potential to stimulate weediness and invasiveness in the invasive weedy *R. sativus* in both ruderal and agrestal environments, regardless of the direction of hybridization. This could promote the introgression and spread of crop alleles -including alleles with resistance to herbicides, insects, pathogens, viruses and tolerance to abiotic stresses (genetically modified or not)- into weed populations, promoting the evolution of novel weed biotypes. We suggest that subsequent long-term full life-cycle experiments in multiple locations are necessary to assess the generality of our results. Further research should also study the phenotypic variation of functional traits on which selection can act, as well as the strength and direction of direct and indirect selection operating on them using phenotypic selection analysis to better understand the process of adaptive evolution that may accompany the evolution of weeds and invasives.

## ACKNOWLEDGMENTS

This work was supported by the Agencia Nacional de Promoción Científica y Tecnológica (ANPCyT) PICT 2017-0473. We thank the National Research Council of Argentina (CONICET) for a fellowship for FH and RBV, and two anonymous reviewers for their helpful feedback that greatly improved the manuscript.

